# Hook-transmitted torque drives spinning and precession of bacterial flagella

**DOI:** 10.64898/2026.05.26.727843

**Authors:** G. Donini, Silvio Bianchi, R. Di Leonardo

## Abstract

Bacterial propulsion is powered by torque generated by the flagellar motor and transmitted to the flagellum through the hook, a short deformable structure acting as a flexible joint. Despite the hook being essential for propulsion, the physical mechanism governing torque transmission remains poorly characterized. Here, we show that the torque consists of two components: one parallel to the flagellum, responsible for rapid spin, and another along the motor axis, which induces precession of the former component. Evidence of such a two-component torque also emerge in bead assay experiments, where the attached bead can exhibit complex trajectories rather than simple circular motion. We introduce a minimal mechanical model that captures both the spin/precession dynamics and the commonly observed circular orbits.

## Introduction

The bacterial flagellar motor is a powerful rotary nanomachine that converts electrochemical energy into mechanical work [6, 19]. This 50 nm protein complex in *Escherichia coli* typically rotates at a frequency of 100 Hz, generating a torque of order 0.5 pN*µ*m. The torque is transmitted to the flagellum through the hook [38], a short (60 nm) elastic filament that acts as a universal joint [7, 17, 18, 22, 39] and is highly conserved across species [50]. The hook enables the mechanical flexibility required for flagella to align with the cell body, forming a coherent bundle [42], which drives propulsion at speeds of about 20 *µ*m/s. While torque generation in the motor is widely studied and modeled [27, 29, 48], much less attention has been devoted to what happens next, specifically, to how the torque is transferred to the flagellum. In this context, the role of the hook is typically disregarded. Some numerical models of microswimmers prescribe a fixed rotational speed for the flagellum along its axis, treating the torque applied at its base as a quantity to be determined [13, 36], while others assume a motor torque parallel to the flagellum axis [16, 49]. In contrast, a different modeling approach assumes the motor torque to be perpendicular to the cell surface [11, 25, 47]. This assumption is also implicitly made in single-motor studies employing the bead assay [8, 27, 29, 33, 41, 45, 46, 52]. Here, using both simple mechanical arguments and observations of the motion of different loads coupled to the motor, we demonstrate that the transmitted torque has two components: one parallel to the flagellum axis, causing rapid spin, and another directed along the motor axis, causing precession. To test our model, we focus on experiments performed using the bead assay [20], by far the most successful technique employed to study the bacterial flagellar motor. In this approach, a nano- or microsphere is coupled to the motor through the hook or the flagellum, following appropriate functionalization. Over the years, this technique has enabled the characterization of several aspects of motor dynamics, including the torque–speed relationship [8, 21, 27, 29, 33, 52], stator binding and unbinding dynamics [26, 32, 35, 37, 40, 44], and motor reversal in chemotaxis [1, 9, 51]. More recently, the bead assay has also played an important role in uncovering aspects of bacterial electrophysiology [2, 5, 23, 24, 28, 43]. Despite these successes, we identify a discrepancy in the literature between the widespread use of the bead assay and the limited understanding of how the motor torque is transmitted to the bead through the hook. We show that beads can exhibit either the simple circular trajectory commonly reported in the literature or a more complex, often disregarded class of “curly” trajectories, characterized by the superposition of spin and precession. Our minimal model captures this rich phenomenology and provides a framework for studying both microswimmer simulations and single-motor dynamics.

## Results

### Motor torque has two components

In this first section, we derive the equations governing torque transmission through the hook and present direct experimental evidence supporting them. Fig. 1(a) shows a closeup sketch view of a cell outer surface. The hook is a universal joint connecting the motor, whose axis is directed along 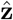, to the flagellum which is considered a rigid rod oriented along 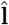. We describe the hook motion as the combination of a rigid rotation about the motor axis 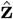 and a spin about its curved axis, with unit tangent 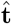 . The local angular velocity is given by 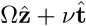, where Ω and *ν* are the angular speeds of the two rotations. At the base of the hook, 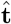 is aligned with 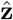, so the angular velocity of the motor is 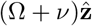 . In contrast, at the tip of the hook 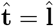, and therefore the angular velocity of the flagellum is:

**Figure 1.**
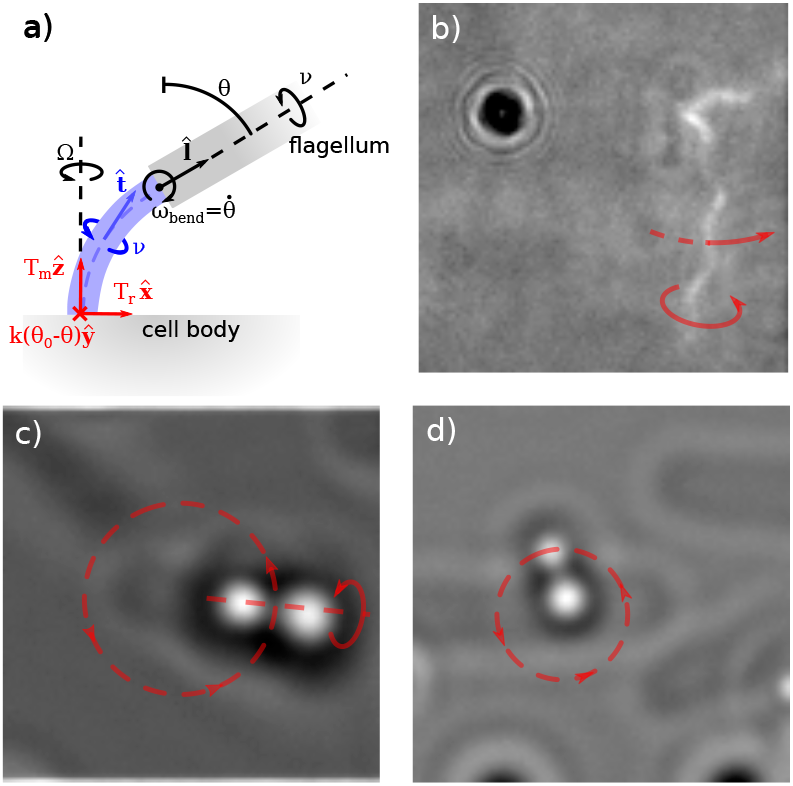
(a) Sketch of the hook connecting the cell body to the flagellum. The hook rotates about the motor axis with angular speed Ω and spins about its local curvilinear axis with angular speed *ν*. Elastic deformations modify the pitch angle *θ* at the hook tip, with angular speed *ω*_bend_. The motor provides a fixed torque *T*_*m*_ along 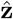 . A reaction component 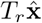 arises when torque is transmitted along the direction 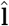 . The elastic torque *T*_el_ = *k*(*θ*_0_ − *θ*) along ŷ accounts for the hook bending response. The resulting torque balance is given in Eq. 3. (b) A cell adhering to a coverglass spins its flagellum about its axis with speed *ν* while simultaneously undergoing precession with speed Ω. (c) Spin–precession mechanism visualized with a flagellum attached to a bead-pair. (d) A short flagellar stub connected to a bead-pair. Contact forces between the bead-pair and the cell body cancel in-plane torque components, allowing pure rigid-body rotation. For (b–d) see also Supplemental Movie 1.

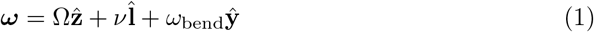

where, for completeness, we include a third component, *ω*_bend_**ŷ**, arising from the elastic bending of the hook, perpendicular to the components along 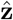 and 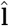 .

Due to the small size of the hook, its hydrodynamic drag coefficients are negligible compared to those of the load attached at its end. Under this assumption, the net torque acting on the hook vanishes; consequently, the torque **T** applied by the motor to the hook is equal to the torque transmitted by the hook to the flagellum. Placing the origin at the flagellar motor, we express the torque exerted by the motor on the hook in its most general form as the sum of three mutually orthogonal components:

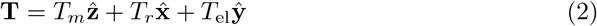

The motor provides a torque *T*_*m*_ along 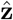 . The component along **ŷ** arises from the elastic bending of the hook; we model it as *T*_el_ = *k*(*θ*_0_ − *θ*) which restores the hook pitch angle *θ* to its equilibrium value *θ*_0_. The component along 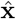, *T*_*r*_, is a reaction torque whose amplitude will be determined below. The power provided by the motor *P*_motor_ must be equal to that dissipated by viscous drag *P*_viscous_ plus the rate at which the elastic energy is stored *P*_elastic_. The three terms are as follows: *(I) P*_motor_ = *T*_*m*_(Ω + *ν*), obtained by multiplying the angular velocity of the hook base, which is directed along 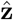, with the *z* component of **T**. *(II) P*_viscous_ = **T** · ***ω*** + **f** · **v**, where **f** is the force provided by the hook, while ***ω*** and **v** are the angular and linear velocities of the load connected at its end. Keeping the origin placed at the motor and for a vanishing hook length (*i*.*e*., the flagellum’s end is directly connected to the origin), we have **v** = 0 simplifying the expression to *P*_viscous_ = **T** · ***ω***. 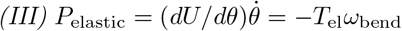 The relation *P*_motor_ = *P*_viscous_ + *P*_elastic_ leads to an expression in which the components of ***ω*** are simplified, resulting in the formula 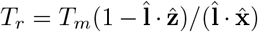 Substituting this into Eq. 2 and carrying out the algebra yields a closed-form expression for **T**:

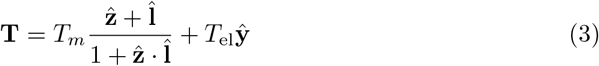

For a step by step derivation, see supplementary materials, section II. Summarizing, in the approximation where the hook size and drag are negligible, the transmitted torque has a component driving the spinning of the flagellum and another one that induces its precession. The elastic contribution simply adds the torque arising from the bending of the hook.

Direct evidence of the above formula is given by fluorescently stained flagella [3]. In Fig. 1(b) we show a cell adhering to a coverglass rotating a flagellum. The flagellum spins rapidly around its axis 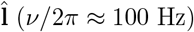 but also rotates around the motor axis 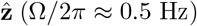 (see also Supplemental Movie 1). Since the motor is located at the top of the cell while the flagellum lies approximately in the *x*-*y* plane, we consider the hook to be bent by 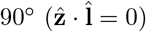 . From fluorescence images, we extract the length, radius, and pitch of the flagellum and, using the Rotne–Prager method, compute the drag coefficients [3, 14] *γ*_‖_ and *γ*_⊥_, associated with rotations about the flagellar axis and the motor axis, respectively. Eq. 3 predicts that the torque components along 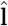 and 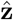 have the same magnitude *T*_*m*_ and are responsible for flagellar spin and precession, respectively. From the measured spinning frequency, we obtain *T*_*m*_ = *γ*_‖_*ν* ≈ 0.5 pN*µ*m, a value comparable to those reported in the literature [4, 10, 12]. Since *T*_*m*_ = *γ*_⊥_Ω also holds, the estimates of *γ*_‖_ and *γ*_⊥_ allow us to obtain an approximate value for the ratio between the precession and spinning frequencies. We estimate Ω*/ν* = *γ*_‖_*/γ*_⊥_ ≈ 0.25*/*100, which is of the same order of magnitude as the measured value, Ω*/ν* ≈ 0.5*/*100. A similar phenomenology is observed in Fig. 1(c), where a bead-pair formed by two irreversibly attached 1 *µ*m polystyrene beads adheres to a sticky flagellum (see also Supplemental Movie 1). The bead-pair spins about its major axis (*ν/*2*π* ≈ 20 Hz) while simultaneously rotating around the motor axis (Ω*/*2*π* ≈ 2 Hz). Again, assuming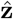 and 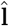 to be orthogonal and computing the drag coefficients of the bead-pair using the Rotne-Prager method, we obtain Ω*/ν* = *γ*_‖_*/γ*_⊥_≈ 2.7*/*20, in good agreement with the observed ratio Ω*/ν* ≈ 2*/*20. This rotation/precession motion is not observed when the load attached to the motor is subject to additional constraints. For example, when the bead-pair is attached to a short flagellar stub, the presence of the cell body imposes a geometrical constraint that allows rotation only around the motor axis, as shown in Fig. 1(d) (see also Supplemental Movie 1). In this case, the forces between the bead-pair and the cell body generate an additional reaction torque that cancels the *x*-*y* components of Eq. 3, resulting in a total torque that is simply 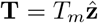 .

### Bead assay revisited

In the bead assay, a nano- or micron-sized bead is attached to a flagellar stub of a cell adhering to a surface. We start by showing the case of a bead with diameter 2*a* = 1.6 *µ*m performing a rigid rotation in Fig. 2(a). In this case, the bead follows a circular trajectory, which in practice corresponds to the two-dimensional projection of a circle, *i*.*e*. an ellipse [53]. Indicating with (*x, y*) the bead position we construct the complex coordinate *ρ*(*t*) = *x*(*t*) + *iy*(*t*) and then compute the modulus of the Fourier transform |*ρ*(*f*) |. For an almost perfect circular trajectory, as the one shown in Fig. 2(a), |*ρ*(*f*) | displays a single sharp peak, although a second harmonic may appear (case not shown). In contrast to these cases, we occasionally observe beads of the same size drawing a curly trajectory, as shown in Fig. 2(b). When this happens, two peaks become visible, with frequencies that are not trivially multiples of each other. A direct qualitative observation of these two behaviors is obtained by using particles that display recognizable features, which allow one to visualize their orientation. Fig. 2(c–d) shows a bead rotating rigidly and a bead performing spin–precession motion, respectively (see also Supplemental Movie 1).

**Figure 2.**
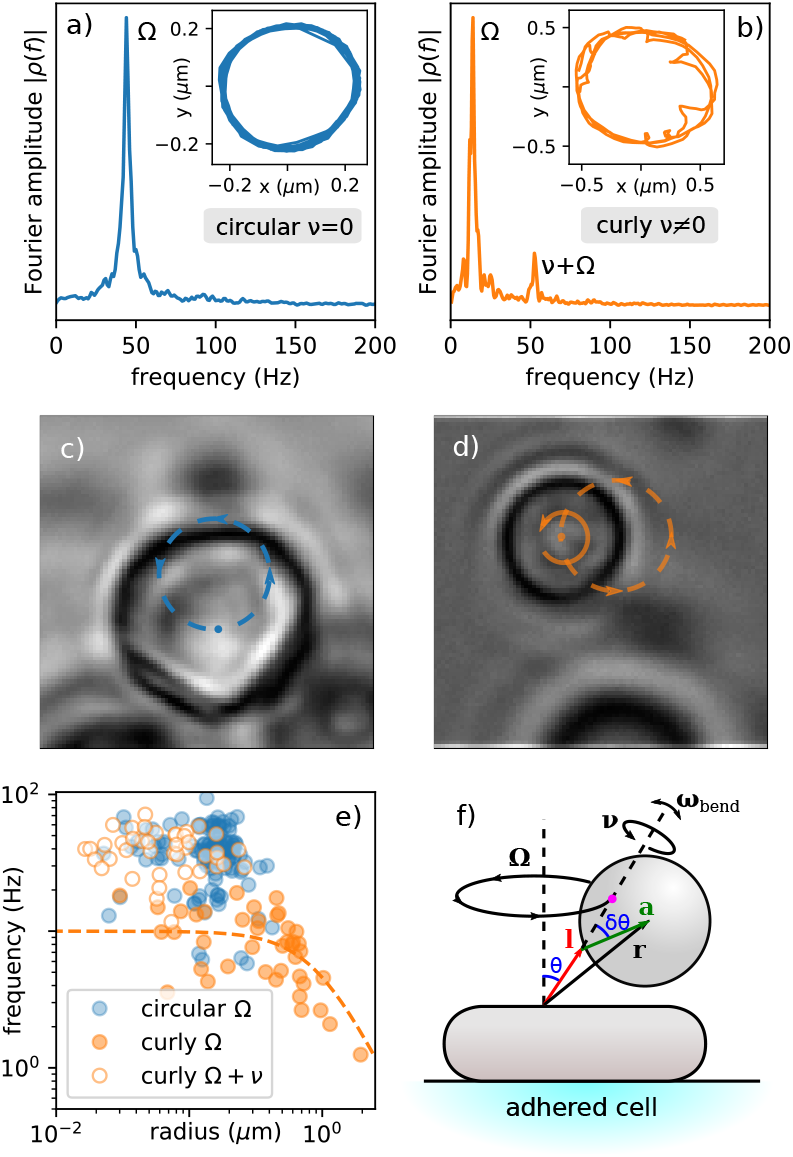
(a-b) Fourier spectra of *ρ* = *x* + *iy* for a rigid circular rotation (a) and a curly trajectory (b). Insets show the corresponding bead trajectories in the *x*-*y* plane. (c–d) Imperfections in some particles allow their rotation about their centers to be visualized. The particle in (c) undergoes rigid-body rotation, whereas the one in (d) exhibits the combined spin/precession motion (see also Supplemental Movie 1). (e) Rotation frequency as a function of trajectory radius for circular trajectories (blue points). For curly trajectories the radii and frequencies, associated with the slow (Ω) and fast (Ω + *ν*) Fourier peaks, are shown as filled orange and open orange points, respectively. The orange dashed line shows the trend Ω ∼ 1*/*(*γ*_*R*_ + γ*r*^2^) (f) Sketch of the bead assay geometry showing the rotation components. The bead position is **r** = **l** + **a**, where **l** is the flagellar stub and **a** connects the attachment point to the bead center. The projection of **r** along **l** (magenta point) rotates around the motor axis 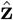 with angular velocity Ω, while the bead additionally spins about **l** with angular velocity *ν*.

The circular trajectory corresponds to a rigid rotation with respect to the motor axis (*ν* = 0). This occurs when the bead touches the cell wall: contact forces arise to cancel the torque components in the *x*-*y* plane and prevent the bead from rotating around the flagellum axis with angular speed Ω. Therefore, the bead experiences a torque 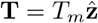 . Circular trajectories can also occur without collisions between the cell and the bead, for a range of geometrical conditions; these cases will be described in the next section.

The curly trajectory corresponds to the case where *ν* ≠ 0 so that rotation around the motor axis is accompanied by a spin of the flagellum. In the Fourier amplitude |*ρ*(*f*) |, the frequency of the two peaks correspond to the angular speeds Ω and Ω + *ν*. To justify that, we refer to Fig. 2(f). For simplicity, we assume that the hook is sufficiently rigid so that 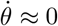, implying 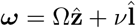 . The bead position is given by **r** = **l** + **a**, where **l** represents the flagellum and **a** is the vector from the attachment point to the bead center. In the simple case where both *θ* and *δθ* are small, rotation about the motor axis produces circular motion in the *x*-*y* plane with radius *a* sin *θ* cos *δθ* + *l* sin *θ* ≈ *aθ* + *lθ* and angular speed Ω. Rotation around the flagellum, which is approximately parallel to the *z* axis, generates an epicyclic motion with radius *aδθ* and angular speed Ω + *ν*. Consequently, the resulting in-plane complex coordinate can be approximated as 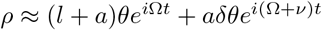 which explains the two peaks observed in Fig. 2(b). In Supplemental Information, we demonstrate that the identification of these two peaks with the angular speeds Ω and Ω + *ν* persists for arbitrary values of *θ* and *δθ*.

From Fourier amplitudes such as the ones shown in Figs. 2(a-b) we obtain the peak frequency *f* ^*^ and its corresponding radius |*ρ*(*f* ^*^)| . Circular trajectories are shown as blue points in Fig. 2(e). For curly trajectories, the high and low frequency peaks are plotted as filled and open orange points, respectively. Points relative to the curly fast peak form a cluster that partially intersects with the one relative to the circular trajectories. Typically, for curly trajectories, the slower Ω peak corresponds to a radius approximately one order of magnitude larger than that associated with the Ω + *ν* peak. For a rotation around the motor axis 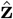 the corresponding drag is *γ*_*R*_ + Γ*r*^2^, with *γ*_*R*_ = 8*πηa*^3^ and Γ = 6*πηa*, where *a* = 0.8 *µ*m is the bead radius, *r* is the orbit radius and *η* is the water viscosity. We therefore expect Ω ∼ 1*/*(*γ*_*R*_ + Γ*r*^2^) a trend that is plotted by orange dashed in Fig. 2(e).

One of the main applications of the bead assay is to provide a straightforward measurement of the motor torque by simply multiplying the rotational speed by the drag coefficient. To our knowledge, cases in which beads exhibit unclear motion with curly trajectories are usually discarded. Curly trajectories appear more frequently when the flagella are not shortened by shear flow. We acquired 180 trajectories using 1.6 *µ*m beads. Among these, 71 belonged to a sample in which the flagella had been damaged by shear flow, following the standard bead assay procedure. The other 109 trajectories corresponded to a sample in which the shear flow treatment was skipped.About 37% of the beads followed a curled trajectory when attached to flagella that had not been sheared, compared to 2.8% in the case of sheared flagella. A possible explanation is that, for shorter flagellar stubs, the bead motion is more likely to be constrained by the nearby cell body, although, as we show in the modeling section, a circular motion can also occur in the absence of any bead–cell collision. As a concluding remark, we point out that for curly trajectories the motor speed Ω + *ν* corresponds to a secondary peak, which is typically smaller or even barely visible, as it is dominated by the peak associated with Ω. This can easily lead to misinterpretation, where the larger peak is incorrectly identified as the motor speed, so that automated analyses may return incorrect estimates of both motor speed and motor torque.

### Simplified model

We consider the sketch in Fig. 2(f). Placing the origin at the motor, the torque transmitted by the hook to the flagellum is given by Eq. 3. If we neglect the drag of the flagellar stub, the same torque **T** is transmitted to the bead. We then shift the origin to the bead center, located at **r** = **l** + **a**. In this reference frame the rotational and translational drag coefficients are those of a sphere, *γ*_*R*_ and Γ respectively, while roto-translational couplings vanish. In the shifted reference frame, the torque must be transformed accordingly. The resulting torque is **T** − **r** × **f**, where **f** is the force exerted by the flagellum on the load. Lastly, we include an external force **f**_ext_ acting on the bead. We consider cases in which, in the reference frame centered at the bead, this force does not result in a torque, as in optical trapping or when the bead interacts with a nearby surface.

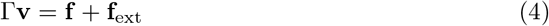

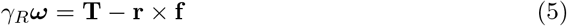

Neglecting the elastic deformations of the flagellar stub, both vectors **a** and **l** undergo rigid-body motion, so that 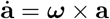 and 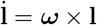 .This in turn implies 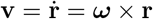 . Eqs. 4-5 simplify to:

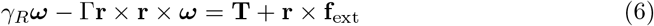

With **T** defined by Eq. 3 and **f**_ext_ being a function of the bead position, Eq. 6 can be easily integrated over time. For now, we focus on the case where no external forces are acting on the bead (**f**_ext_ = 0), meaning we are in a situation where the bead never contacts the cell body surface. Fig. 3(a-c) shows a comparison between three sample experimental curly trajectories of 1.6 *µ*m beads displaying qualitatively different patterns. These trajectories are compared with numerically integrated ones in Fig. 3(d-f). Drag coefficients Γ and *γ*_*R*_ used in the simlutaions are those of a 1.6 *µ*m bead in bulk fluid. The hook torsional elastic constant is chosen as *k* = 3 pN*µ*m/rad, a value that is in the range of that reported in the literature [31, 53]. We vary the flagellum length *l*, the hook’s equilibrium pitch angle at rest *θ*_0_, the angle *δθ* between **l** and **a**, and the torque *T*_*m*_ to reproduce qualitatively the behavior observed experimentally. These values are *θ*_0_=[6^°^, 29^°^, 37^°^], *δθ*=[6^°^, 11^°^, 11^°^], C*l*=[0.2, 0.1, 1.5] *µ*m and *T*_*m*_=[3, 4, 3.2] pN*µ*m in Fig. 3(d), (e) and (f), respectively. In the simulations shown in Fig. 3(d–f), the bead never comes into contact with the cell body, modeled as a spherocylinder of diameter 0.8 *µ*m. The condition **f**_ext_ = 0 in Eq. 6 therefore remains valid.

**Figure 3.**
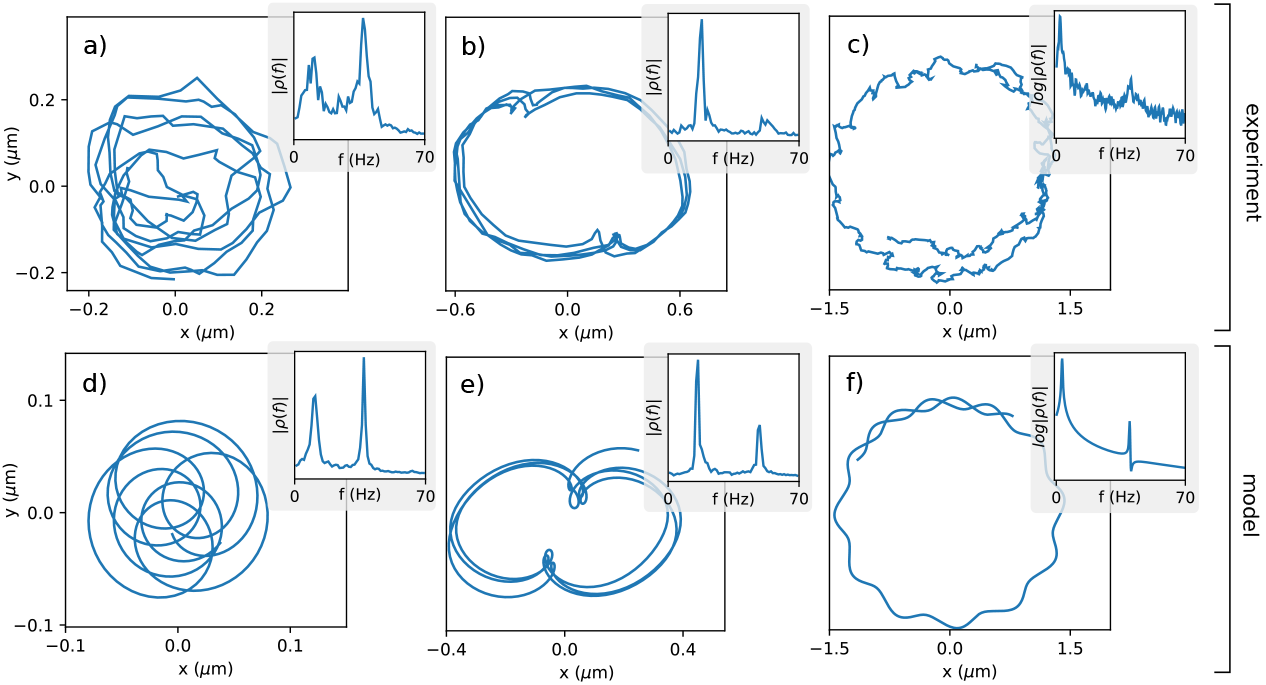
(a-c) Sample experimental curly trajectories along with their Fourier amplitude. (d-f) Numerically generated trajectories using Eq. 3 and Eq. 6 with **f**_ext_ = 0. Parameters in the simulation are adjusted to reproduce the qualitative behavior of the experimental trajectories.

So far, Eq. 6 has been shown to generally yield a solution for ***ω*** containing both Ω and *ν* components, such that the bead undergoes a spin/precession motion, giving rise to a curly trajectory. However, for a range of parameters, Eq. 6 results in a bead performing a circular trajectory. These solutions are those for which ***ω*** is such that *ν* = 0. The component *ν* can be recovered unambiguously from ***ω***. In Eq. 1, we take the scalar product with 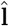 or 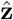 to obtain two equations that can be solved for Ω and *ν*:

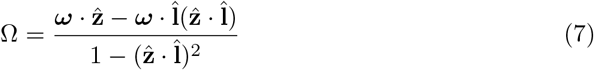

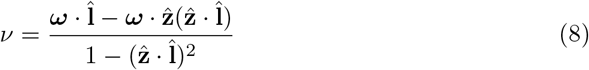

Following Eq. 8 we have that *ν* = 0 when 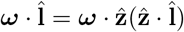 or more simply when *ω*_*x*_*l*_*x*_ + *w*_*y*_*l*_*y*_ = 0. Fig. 4(a) shows a simulation of a 1.6 *µ*m bead with parameters *T*_*m*_ = 1 pN*µ*m, *θ*_0_ = 27^°^ *δθ* = 17^°^ and *l* = 0.1 *µ*m.

**Figure 4.**
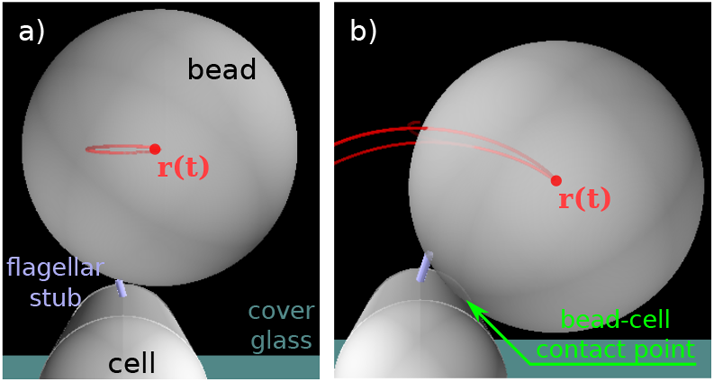
(a) A circular trajectory can be obtained integrating Eq. 6 even when the bead never touches the cell body (**f**_ext_ = 0). (b) When the bead is in contact with the cell, steric forces arise (**f**_ext_ 0) that hinder the rotation about the flagellum axis (*ν* = 0). The bead slides along the cell body surface resulting in a smooth trajectory (see also Supplemental Movie 2).

A trajectory having a single well-defined frequency is also achieved when the bead touches the cell body. We include external forces **f**_ext_ ≠ 0 representing the contact repulsion between the cell body and the bead. Fig. 4(b) shows the simulation with parameters *T*_*m*_ = 1 pN*µ*m, *θ*_0_ = 45^°^ *δθ* = 45^°^ and *l* = 0.2 *µ*m. Rotation along the flagellum axis is impeded by the steric forces between the cell body and the bead.

### Trapping does not block flagellum spin

We show here additional phenomenology associated with the spin/precession rotation induced by the hook. To do that, we observe bead motion while constraining selected degrees of freedom. With both angular clamps, such as magnetic tweezers [30, 32, 45, 46], and torque clamps [15], the motor rotation is effectively constrained. Differently, optical tweezers directly control the bead position while still allowing rotations about the bead center due to the bead’s spherical symmetry and the elasticity of the hook [34]. A sketch of the experiment is shown in Fig. 5(a). Fig. 5(b) shows the *x*–*y* trajectory of a bead at different optical trap powers. At low laser power, the bead is weakly confined and effectively undergoes free motion around the motor center. As the laser power is increased, the trajectory contracts and its center shifts toward the optical trap position. In an *x*–*y* reference frame centered on the motor, the average angular position *ϕ* is obtained by equating the motor torque along *z* to that due to optical confining, *k*_trap_(*ϕ*_trap_ − *ϕ*). It follows that:

**Figure 5.**
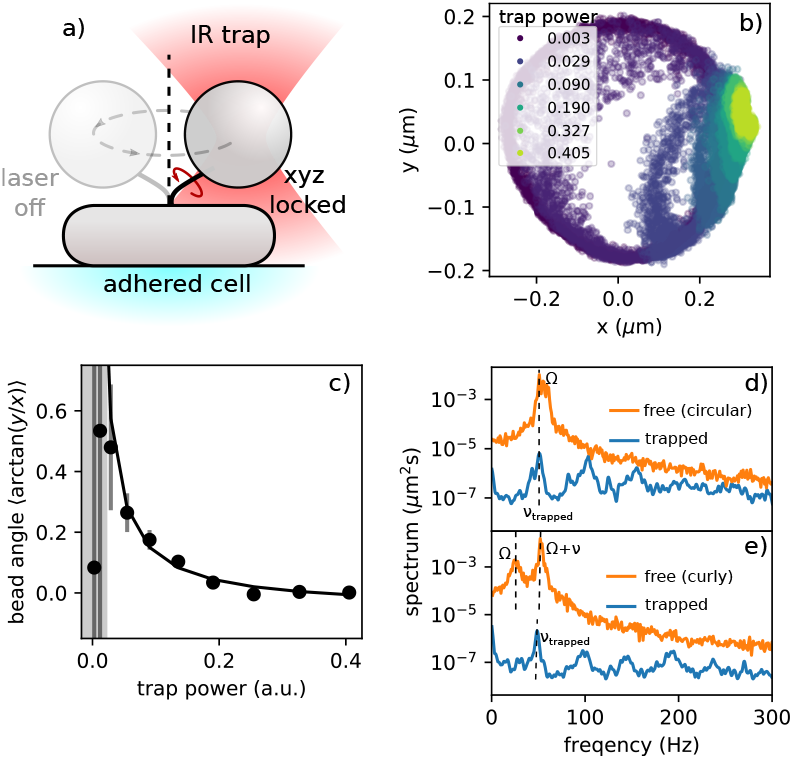
(a) A bead rotating around the motor axis is still able to rotate when its position is confined by an optical trap. (b) Bead position for different trapping laser power. (c) Mean polar angle of the *x*-*y* position as a function of trapping power. Points in the shaded area correspond to power levels that are too low to prevent the bead from orbiting around the motor axis. (d–e) Comparison of the Fourier amplitude of *ρ* = *x* + *iy* between freely rotating beads and trapped beads. Both when the free bead rotates rigidly (d) and when displaying a curly trajectory (e), small oscillations due to the hook spin are visible.

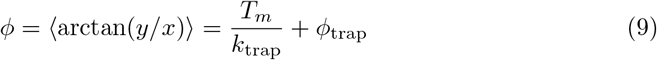

where *x* and *y* are the bead coordinates, *ϕ*_trap_ is the angular position of the trap, and *k*_trap_ is the trap stiffness, which scales linearly with trapping power. Consequently, the average bead position approaches the trap focus as the laser power increases, as shown in Fig. 5(c). At the highest trapping power, *P* = 0.405 (a.u.), the bead is tightly confined, yet it still exhibits a clear spinning motion. This is evident from the Fourier amplitude, |*ρ*(*f*) |, which displays a fundamental peak at a frequency very similar to that observed in the absence of the optical trap (free bead), as shown in Fig. 5(d).

For the trapped bead, the peak in |*ρ*(*f*)| corresponds to a rotation around 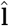 . According to Eq. 3, the torque projection along this direction is *T*_*m*_, resulting in a rotational speed *ν*_trapped_ = *T*_*m*_*/γ*_*R*_. A similar value is obtained for the free bead. Since the motion is circular, there is no rotation along 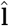, so we consider the torque projection along 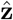, which is again *T*_*m*_ according to Eq. 3. Denoting the orbit radius around the motor axis as *r*, the drag is *γ*_*R*_ + Γ*r*^2^. With *r* ≈ 200 nm and *a* = 0.8 *µ*m we can approximate *γ*_*R*_ ≫ Γ*r*^2^ and thus obtain a rotational speed that is approximately equal to that obtained in the trapped case, *i*.*e*. Ω = *T*_*m*_*/γ*_*R*_. Fig. 5(e) shows a similar comparison with for a bead performing spin/precession motion. When trapped, the peak corresponding to Ω disappears, while the second peak slightly decreases in frequency. To provide a rough explanation, we consider Eq. 6 and, for simplicity, neglect the drag contribution proportional to Γ, as done previously. With no external forces acting on the bead, we have *γ*_*R*_***ω*** = **T**, which, according to Eqs. 7-8, gives both Ω and *ν* equal to 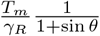 Therefore 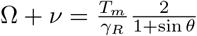 is slightly higher than the speed of the trapped bead that is *ν*_trapped_ = *T*_*m*_*/γ*_*R*_ as in the case of Fig. 5(e).

In Supplemental Movie 1, the particles shown in Fig. 2(c–d) are observed both while moving freely and while trapped. Surface features on the beads allow us to visualize that, in both cases, whenever a bead follows a circular or curly trajectory while free, it spins while held in position by an optical trap, providing a direct visualization of the behavior shown in Fig. 5(d–e).

## 1 Conclusions

Direct observations show that flagellum rotation can be decomposed into a spin about its own axis and a precession around the motor axis. This behavior, which persists even when the flagellar hook is bent by approximately 90^°^ as in Fig. 1(a-b), can be explained only by a torque composed of a constant component along the motor axis and a component along the flagellum axis undergoing precession.

In the bead-assay context, the same spin/precession motion is present, although it is often suppressed for certain geometrical configurations or hindered by the presence of the cell body (see Fig. 4). When the rotation is not rigid-like, the motor speed is Ω + *ν*, which is often associated with a smaller or barely visible peak in the spectrum. A significant error may arise if the frequency corresponding to the peak at Ω, as in the case of Fig. 2(b), is mistaken for the motor speed. This description is supported by the analytical model introduced in the previous section. Although simple, the model, when integrated numerically, qualitatively reproduces the non-trivial traces shown in Fig. 3.

In conclusion, we provide experimental and computational evidence supporting the proposed hook torque transmission mechanism. This insight is important for the interpretation of future experimental data, including bead-assay measurements, and for the accurate simulation of microswimmers, particularly peritrichous ones.

## Supplementary Material

### Supplemental Information

Supplemental information to paper

### Supplemental Movie 1

A collection of videos showing both spin/precession motion and rigid rotation when the motor is connected to various loads.

### Supplemental Movie 2

Numerical simulations of a bead assay reproducing spin/precession motion and rigid rotation.

## Acknowledgments

This project has received funding from the European Research Council (ERC) under the European Union’s Horizon 2020 research and innovation programme (Grant Agreement No. 834615).

