## Supplementary material for "Hook-transmitted torque drives spinning and precession of bacterial flagella": Supplmental Information

(Dated: May 26, 2026)

### CONTENTS

I. Sample preparation 2

II. Derivation of Eq.3 3

III. Fourier Peaks of bead trajectory 4

References 6

### I. SAMPLE PREPARATION

We used the *E. coli* strain PL4 (HCB1826) [1], which has a deletion of the cheY gene, resulting in a lack of the tumbling mechanism in the cells. This strain expresses a "sticky" mutation of the FliC protein, so that a flagellar stub adheres to a microsphere through hydrophobic interaction. Bacteria from frozen glycerol stock were streaked on a Petri dish containing 1.5% agar and lysogeny broth (LB: 1% tryptone, 0.5% yeast extract and 0.5% NaCl). A single colony was inoculated into LB and grown overnight in a shaking incubator at 30 °C and 150 rpm. The overnight culture is diluted 100-fold into 5 mL of the previous medium, grown at 30°C, 150 rpm. At OD≈0.6, cells are collected by centrifugation (1300 rcf, 5'). The resulting pellet is washed twice by centrifugation (1300 rcf, 5') with deionized water buffer, with 66 mM NaCl, 1 mM photassium phosphate (pH 7.0) and 0.1 mM EDTA. The flagellar filaments are sheared by passing the bacteria back and forth 60 times between two syringes (23-gauge) joined by a thin piece of polyethylene tubing. The bacterial concentration is finally adjusted to the working OD≈3.

The sample is prepared as follows: we create a small chamber that we fill with poly-L-lysine (0.01% in deionized water). After 5 minutes we wash the chamber with motility buffer, using some blotting paper to create a flux. We then add bacteria and wait a few minutes for a layer of cells to adhere to the glass, then we wash again with motility buffer and add the polystyrene beads (diluted in deionized water, 0.1% volume fraction). After a few minutes, when it is possible to see a good number of beads rotated by motors, we flush the diffusing ones. In addition to single beads, bead-pairs as the ones shown in Fig. 1(c-d)

*

of main text are found occasionally. Beads exhibiting surface imperfections, such as those shown in Fig. 2(c-d) of main text, are polystyrene beads that were deposited onto a glass substrate and coated with a 100 nm layer of chromium. After sonication and resuspension in water, the microspheres still exhibit small chromium patches.

Preparation of the cells with fluorescently labeled flagella in Fig. 1(b) of main text has been described in [2]. Briefly, the strain is derivated of HCB1737 $\Delta cheY$ [3], a smooth swimmer that expresses a modified flagellin (fliC) that allows filaments to be specifically labeled with the maleimide fluorescent dye Alexa Fluor 680 $C_2$ Maleimide (A-20344; Life Technologies, Carlsbad, California). The strain is also photokinetic by deleting the atp operon and transforming the strain with a plasmid expressing the protein proteorhodopsin, that acts as a light inducible proton pump. In the absence of oxygen and nutrients, the proto motive force, which powers the flagellar motors, is mainly generated by proteorhodopsin [4–6]

### II. DERIVATION OF EQ.3

We present here a step-by-step derivation of Equation 3 of the main text. We recall the definitions of $\boldsymbol{\omega}$ and $\mathbf{T}$:

$$\boldsymbol{\omega} = \Omega\hat{\mathbf{z}} + \nu\hat{\mathbf{l}} + \omega_{\text{bend}}\hat{\mathbf{y}} \tag{1}$$

$$\mathbf{T} = T_m\hat{\mathbf{z}} + T_r\hat{\mathbf{x}} + T_{\text{el}}\hat{\mathbf{y}} \tag{2}$$

In the main text, we provided the expressions for the power provided by the motor at the hook base $P_{\text{motor}}$, the power dissipated by the viscous drag of the load $P_{\text{viscous}}$, and the rate at which elastic energy is stored $P_{\text{elastic}}$:

$$\begin{aligned} P_{\text{motor}} &= T_m\left(\Omega + \nu\right) \\ P_{\text{viscous}} &= \mathbf{T}\cdot\boldsymbol{\omega} \\ P_{\text{elastic}} &= -T_{\text{el}}\omega_{\text{bend}} \end{aligned}$$

Energy conservation gives the relation $P_{\text{motor}} = P_{\text{viscous}} + P_{\text{elastic}}$. Using Eqs. 1 and 2, we list below the algebraic simplification that leads to the expression for $T_r$:

$$
\begin{aligned}
P_{\text{motor}} &= P_{\text{viscous}} + P_{\text{elastic}} \\
T_m\left(\Omega+\nu\right) &= \mathbf{T}\cdot\boldsymbol{\omega} - T_{\text{el}}\omega_{\text{bend}} \\
T_m\left(\cancel{\Omega}+\nu\right) &= \cancel{T_m\Omega} + T_m\nu\ \hat{\mathbf{z}}\cdot\hat{\mathbf{l}} + T_r\nu\ \hat{\mathbf{x}}\cdot\hat{\mathbf{l}} + \cancel{T_{\text{el}}\omega_{\text{bend}}} - \cancel{T_{\text{el}}\omega_{\text{bend}}} \\
T_m\cancel{\nu}\left(1-\hat{\mathbf{z}}\cdot\hat{\mathbf{l}}\right) &= T_r\cancel{\nu}\left(\hat{\mathbf{x}}\cdot\hat{\mathbf{l}}\right)
\end{aligned}
$$

To obtain the concise expression presented in Equation 3 of the main text, we substitute $T_r = T_m(1-\hat{\mathbf{z}}\cdot\hat{\mathbf{l}})/(\hat{\mathbf{x}}\cdot\hat{\mathbf{l}})$ into the definition of $\mathbf{T}$ in Equation 2:

$$
\mathbf{T} = T_{\text{el}}\hat{\mathbf{y}} + T_m\hat{\mathbf{z}} + T_m\frac{1-l_z}{l_x}\hat{\mathbf{x}} \tag{3}
$$

where $l_x = \hat{\mathbf{x}}\cdot\hat{\mathbf{l}}$, $l_z = \hat{\mathbf{z}}\cdot\hat{\mathbf{l}}$, and thus $l = l_x\hat{\mathbf{x}} + l_z\hat{\mathbf{z}}$. Note that, without loss of generality, we choose our axes such that $\hat{\mathbf{y}}\cdot\hat{\mathbf{l}} = 0$. We then substitute the versor $\hat{\mathbf{x}} = (\hat{\mathbf{l}} - l_z\hat{\mathbf{z}})/l_x$ into Eq. 3:

$$
\begin{aligned}
\mathbf{T} &= T_{\text{el}}\hat{\mathbf{y}} + T_m\left(\hat{\mathbf{z}} + \frac{1-l_z}{l_x}\frac{\hat{\mathbf{l}}-l_z\hat{\mathbf{z}}}{l_x}\right) = \\
&= T_{\text{el}}\hat{\mathbf{y}} + T_m\left(\frac{l_x^2 - l_z + l_z^2}{l_x^2}\hat{\mathbf{z}} + \frac{1-l_z}{l_x^2}\hat{\mathbf{l}}\right)
\end{aligned}
$$

Finally, we use the relation $l_x^2 + l_z^2 = 1$ to obtain the result:

$$
\begin{aligned}
\mathbf{T} &= T_{\text{el}}\hat{\mathbf{y}} + T_m\frac{1-l_z}{1-l_z^2}(\hat{\mathbf{z}}+\hat{\mathbf{l}}) = \\
&= T_{\text{el}}\hat{\mathbf{y}} + T_m\frac{1}{1+l_z}(\hat{\mathbf{z}}+\hat{\mathbf{l}})
\end{aligned}
$$

### III. FOURIER PEAKS OF BEAD TRAJECTORY

We refer to Fig. 1(a) and compute the trajectory of the bead in the ideal case where the hook rigidity is large enough to neglect bending $\omega_{\text{bend}} = \dot{\theta} = 0$. Also in the present case we ignore collisions between the bead and the cell body. For $\theta = 0$ and $\Omega = 0$ we have that the flagellum is parallel with the $z$ axis $\mathbf{l}' = l[0,0,1]$ and

$$
\mathbf{a}' = a\begin{bmatrix}\sin\delta\theta\cos(\nu t) \\ \sin\delta\theta\sin(\nu t) \\ \cos\delta\theta\end{bmatrix}
$$

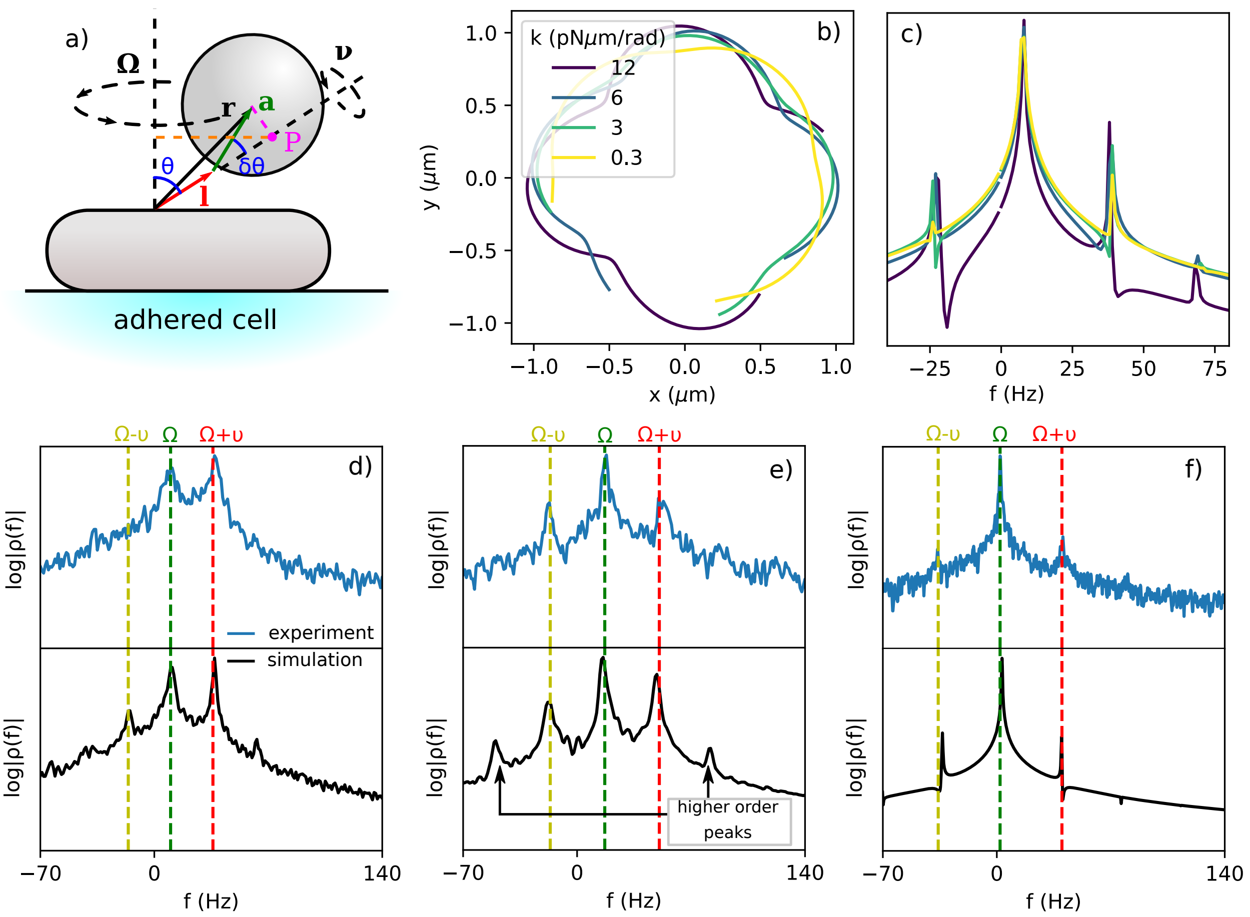


FIG. 1. a) Sketch of bead assay. For $\dot{\theta} = 0$, the Point P rotates around the $z$ axis with speed $\Omega$ and radius $C_\Omega$ indicated by orange dashed line. Magenta dashed line is related with the amplitude of the $\Omega \pm \nu$ peaks with its length being $C_{\Omega+\nu} + C_{\Omega-\nu}$. b) Simulated trajectory for various values of hook rigidity $k$. c) Fourier amplitudes $|\rho(f)|$ of the trajectories shown in (b). d-f) Fourier amplitudes shown in Fig. 3 of main text plotted of over a larger range of frequencies. Higher order harmonics appear when $\dot{\theta} \neq 0$. These are generally suppressed in experimental data, probably because of Brownian noise.

To obtain the general case, with $\theta \neq 0$ and $\Omega \neq 0$, we rotate both vectors by $\theta$ along the $y$ axis and then rotate by an angle $\Omega t$ along the $z$ axis. For $\mathbf{l}$ we obtain:

$$\mathbf{l} = R_z(\Omega t) R_y(\theta) \mathbf{l}' = \begin{bmatrix} \cos(\Omega t) & -\sin(\Omega t) & 0 \\ \sin(\Omega t) & \cos(\Omega t) & 0 \\ 0 & 0 & 1 \end{bmatrix} \begin{bmatrix} \cos\theta & 0 & \sin\theta \\ 0 & 1 & 0 \\ -\sin\theta & 0 & \cos\theta \end{bmatrix} \begin{bmatrix} 0 \\ 0 \\ l \end{bmatrix} = l \begin{bmatrix} \sin\theta\cos(\Omega t) \\ \sin\theta\sin(\Omega t) \\ \cos\theta \end{bmatrix}$$

Similarly for $\mathbf{a}$:

$$\mathbf{a} = R_z(\Omega t) R_y(\theta)\mathbf{a}'$$

We compute the bead position $\mathbf{r} = \mathbf{l} + \mathbf{a}$ and consider only the observable $x$ and $y$. The in-plane complex coordinate $\rho = x + iy$ can be simplified to:

$$\rho = C_\Omega e^{i\Omega t} + C_{\Omega+\nu} e^{i(\Omega+\nu)t} + C_{\Omega-\nu} e^{i(\Omega-\nu)t} \tag{4}$$

with

$$C_\Omega = l\sin\theta + a\sin\theta\cos\delta\theta \tag{5}$$

$$C_{\Omega+\nu} = \frac{a}{2}(1+\cos\theta)\sin\delta\theta \tag{6}$$

$$C_{\Omega-\nu} = \frac{a}{2}(1-\cos\theta)\sin\delta\theta \tag{7}$$

The Fourier amplitude has three peaks. Note that the peak at $\Omega - \nu$ vanishes for small $\theta$. To correctly interpret the amplitudes of the three peaks, we note that the point $P$ in Fig. 1(a) traces a circle in the $x$-$y$ plane with angular speed $\Omega$ and radius $C_\Omega$. The bead center rotates around the axis $\mathbf{l}$ with speed $\nu$ and rotation radius $C_{\Omega+\nu} + C_{\Omega-\nu}$.

When the elasticity of the hook is included, $\dot{\theta} \neq 0$, which introduces additional oscillations in $\mathbf{r}$. Fig. 1(b) shows a simulation of a 1.6 $\mu$m bead with $L = 0.5$ $\mu$m, $\theta = 45°$, $\delta\theta = 25°$, and hook torsional stiffness $k$ taking the values $k = 12, 6, 3, 0.3$ pN, $\mu$m/rad. The corresponding Fourier amplitudes of $\rho$ are shown in Fig. 1(c).

The three peaks in Eq. 4 also appear in the experimental data. Figures 1(d–f) show the same Fourier amplitudes as in Fig. 4 of the main text, plotted over a larger frequency range. Higher-order harmonics are generally suppressed in the experimental data, likely due to Brownian noise. Note also that the data shown in (d), reproduced by simulating a hook with a small angle $\theta \approx 6°$, exhibits a disappearing peak at $\Omega - \nu$, as expected from Eq. 7.

[1] P. P. Lele, B. G. Hosu, and H. C. Berg, Dynamics of mechanosensing in the bacterial flagellar motor, Proceedings of the National Academy of Sciences **110**, 11839 (2013).

[2] S. Bianchi, F. Saglimbeni, G. Frangipane, M. Cannarsa, and R. Di Leonardo, Light-driven

flagella elucidate the role of hook and cell body kinematics in bundle formation, PRX Life **1**, 013016 (2023).

[3] S. Bianchi, F. Saglimbeni, G. Frangipane, and R. Di Leonardo, Flagellar elasticity and the multiple swimming modes of interfacial bacteria, Physical Review Research **4**, L022044 (2022).

[4] J. Arlt, V. A. Martinez, A. Dawson, T. Pilizota, and W. C. Poon, Painting with light-powered bacteria, Nature communications **9**, 768 (2018).

[5] G. Frangipane, D. Dell'Arciprete, S. Petracchini, C. Maggi, F. Saglimbeni, S. Bianchi, G. Vizsnyiczai, M. L. Bernardini, and R. Di Leonardo, Dynamic density shaping of photokinetic e. coli, Elife **7**, e36608 (2018).

[6] S. Bianchi, G. Donini, M. C. Cannarsa, G. Frangipane, and R. Di Leonardo, Dynamic velocity response of e. coli powered by proteorhodopsin, Biophysical Reports (2026).
